# A corollary discharge mediates saccade related inhibition of single units in mnemonic structures of the human brain

**DOI:** 10.1101/2021.07.12.450085

**Authors:** Chaim N. Katz, Andrea G.P. Schjetnan, Kramay Patel, Victoria Barkley, Kari Hoffman, Suneil K. Kalia, Katherine Duncan, Taufik A. Valiante

## Abstract

Despite the critical link between visual exploration and memory, little is known about how single-unit activity (SUA) in the human mesial temporal lobe (MTL) is modulated by saccadic eye movements (SEMs). Here we characterize SEM associated SUA modulations, unit-by-unit, and contrast them to image onset, and to occipital lobe SUA. We reveal evidence for a corollary discharge (CD)-like modulatory signal that accompanies SEMs, inhibiting/exciting a unique population of broad/narrow spiking units, respectively, before and during SEMs, and with directional selectivity. These findings comport well with the timing, directional nature, and inhibitory circuit implementation of a CD. Additionally, by linking SUA to event-related potentials (ERPs), which are directionally modulated following SEMs, we recontextualize the ERP associated with SEM as a proxy for both the strength of inhibition and saccade direction, providing a mechanistic underpinning for the more commonly recorded SEM-related ERP in the human brain.

## Introduction

Humans rely on vision to understand their environment [1], [2], and their primary tool for exploration is the saccade [3], [4] – a ballistic movement of the eyes from one fixation point to another. Given saccades importance, it may seem intuitive that they are inextricably linked to memory [5]–[12] and are associated with prominent electrophysiological responses [13]–[15] in memory–related, mesial temporal lobe (MTL) structures. Anatomical connections between MTL and oculomotor systems have been investigated[9]. However, the cellular mechanisms through which oculomotor structures influence the human MTL remain largely unknown, retarding the understanding of *how* eye movements interact with the memory system. In vision, saccade-related motor signals, in the form of a corollary discharge (CD), prepare the brain for the sensory consequences of a planned movement [16], [17]. Our and others’ recent local field potential (LFP) recordings provide preliminary clues that saccade-related responses in the human MTL may also reflect a CD-like signal. Specifically, we previously demonstrated that saccade-related MTL ERPs are unlikely to reflect visual exafference; visual and saccade-related activity displayed different oscillatory profiles, and saccade-related phase resetting was time-locked to the onset of the motor (saccade) initiation and not the onset of new visual information (fixation) [15]. Moreover, saccadic modulation of MTL electrophysiology has been shown to persist in the dark [18]–[20], again, suggesting that the MTL modulation likely arises from internally generated signals, like a CD. However, a more direct assessment of the CD hypothesis requires recording individual neurons within the human MTL to assess core properties of a CD-like mechanism underlying the saccadic eye movement(SEM) related MTL modulations.

In the most rudimentary implementation of a CD, a copy of the initiating motor command suppresses the sensation produced by the intended motor action [21]–[23]. In this implementation, the circuit motif consists of axon collaterals of primary motor neurons synapsing onto inhibitory interneurons, which inhibit sensory neurons during the movement [16], [17]. Increasingly complex variations on this motif have been presented for the CD-like discharges across the phylogenetic tree; however, inhibition remains a key mechanism by which CDs mediate their effects [17]. Indeed, within the taxonomy for CDs proposed by Crapse and Sommer, the saccadic inhibition/suppression of visual cortical excitability [24] comports well with the inhibition and sensory filtering function of a lower-order CD. This saccadic suppression is considered a requirement for registering a stable visual percept of the world by blocking visual input during ballistic eye movements (although alternative hypotheses exist for the observed saccadic suppression [25]).

Additionally, CDs are thought to help produce a stable visual percept by dynamically shifting receptive fields in anticipation of ensuing eye movements (for a full review, see [26]). Thus, CDs appear to contain information about the direction and magnitude of ensuing saccades. Such a higher-order CD facilitates the transformation of the motor signal to visual coordinates, necessary for anticipatory adjustments of neuronal receptive fields and updating spatial locations [21]. Such a function is well exemplified by its disruption; inhibiting the mediodorsal nucleus, a known node in a higher-order CD circuit, degraded non-human primate’s (NHP) conscious perception of a target shift [27].

In NHPs, CD’s have been proposed as the source of directional information in the entorhinal cortex (EC) [28] and to play a role in spatial mapping within EC [28], [29]. Further research is needed to evaluate the CD hypothesis within the human brain, since only circumstantial LFP evidence exists for a CD mediating SEM related MTL modulations in humans [14], [15], [30]. Thus, despite LFP support for the presence of a CD within MTL structures, such a hypothesis remains wanting of support at the cellular level [31]. Based on the CD literature [17], [24], [27], [31], [32], we propose a framework for amassing evidence of a CD in the human MTL. First, since CDs reflect the anticipatory modulation of sensory and/or higher-order structures, an SEM-related CD in the MTL must modulate firing rates before or during the saccade. Second, since most rudimentary CD circuits involve a prominent inhibition of sensory input, CD-related modulation of MTL activity should be largely inhibitory, evidenced by a decreased firing rate of putative pyramidal neurons and/or increased firing rate of putative inhibitory neurons. Third, there is extensive evidence of the modulation of single unit activity (SUA) in the MTL to visual stimuli [33]–[37]. Thus, any CD-related modulation of activity in the MTL should be distinct from modulation associated with visual input. Finally, as part of an oculomotor circuit, CDs should represent the directional parameters of the ensuing movement. Thus, the CD-related modulation of SUA within the MTL must demonstrate directional tuning (i.e., dissociable neuronal responses to ipsiversive and contraversive saccades).

Within this framework, we leverage the unique opportunity of recording SUA from epilepsy patients undergoing diagnostic stereo-electroencephalography (sEEG). While the participants visually searched natural scene images, we simultaneously recorded SUA from the hippocampus and related MTL structures, along with eye movements. We utilized these recordings to characterize peri-saccadic SUA according to our four criteria, both within the MTL and serendipitously obtained recordings from a control region, the occipital lobes, in two patients. Based on the above criteria, and recognizing the limitations of human-related research, we provide evidence of an SEM-related CD that modulates SUA in the human hippocampus and other related MTL structures.

## Methods

### Participants

Eleven individuals with medically refractory epilepsy participated in this study (Table 1). As part of their clinical assessment, sEEG electrodes (Ad-Tech Medical, WI, USA) were implanted in clinically-determined sites to assess their candidacy for surgical resection of an epileptogenic focus. If patients provided written research consent to microwire recordings, then the sEEG macro-electrodes were supplemented with microwires (Ad-Tech Medical, WI, USA). Microwire implants and all recordings were approved by the Research Ethics Board of the University Health Network.

**Table 1.** Patient summary table. Data for each participant is shown in the respective rows. HPC: hippocampus; AM: Amygdala; PHG: Parahippocampal Gyrus; OCC: Occipital areas

| ID | Sex | Age | Sessions | # Saccades | HPC units | AM units | PHG units | OCC units | Total units |
| --- | --- | --- | --- | --- | --- | --- | --- | --- | --- |
| P89 | F | 45 | 1 | 538 | 13 | 14 | 30 | 0 | 57 |
| P90 | M | 20 | 1 | 414 | 11 | 22 | 0 | 98 | 131 |
| P91 | F | 59 | 1 | 463 | 2 | 3 | 0 | 0 | 5 |
| P100 | F | 19 | 1 | 527 | 25 | 1 | 0 | 0 | 26 |
| P101 | F | 25 | 1 | 625 | 32 | 6 | 10 | 0 | 48 |
| P103 | M | 49 | 1 | 558 | 3 | 0 | 0 | 0 | 3 |
| P107 | F | 64 | 1 | 641 | 0 | 9 | 1 | 0 | 10 |
| P109 | M | 28 | 1 | 593 | 36 | 8 | 0 | 0 | 44 |
| P116 | M | 28 | 1 | 555 | 21 | 58 | 0 | 0 | 79 |
| P125 | M | 24 | 2 | 628 | 14 | 43 | 63 | 21 | 141 |
|  |  |  |  | 821 | 8 | 22 | 22 | 6 | 58 |
| P126 | M | 25 | 1 | 589 | 45 | 0 | 2 | 0 | 47 |
| Total |  |  |  |  | 210 | 186 | 128 | 125 | 649 |

### Experimental Design

The terms “neuron” and “unit” are used interchangeably. To characterize SUA in the time interval surrounding SEMs, we developed a task that involved image presentation and target search (Figure 1A) to engage participants in generating SEMs. The encoding session was divided into 40 trials, each consisting of a fixation cross (1 second), followed by presenting a scene image. Each scene contained four copies of one of two unrelated objects, i.e., targets, that were unrelated to the scene. Participants were instructed to view the individual, serially-presented scene and find as many targets within the scene as possible. If they found all four targets, participants were instructed to continue to move their eyes through the scene to remember and associate the target with the scene. Each scene image was presented for four seconds, after which participants were asked how many targets they found. The trials were grouped into blocks of five trials, with each block having the same embedded target. Following the encoding session, participants were presented with instructions on the ensuing retrieval session. The data presented here is based solely on eye movements during the encoding session.

**Figure 1.**
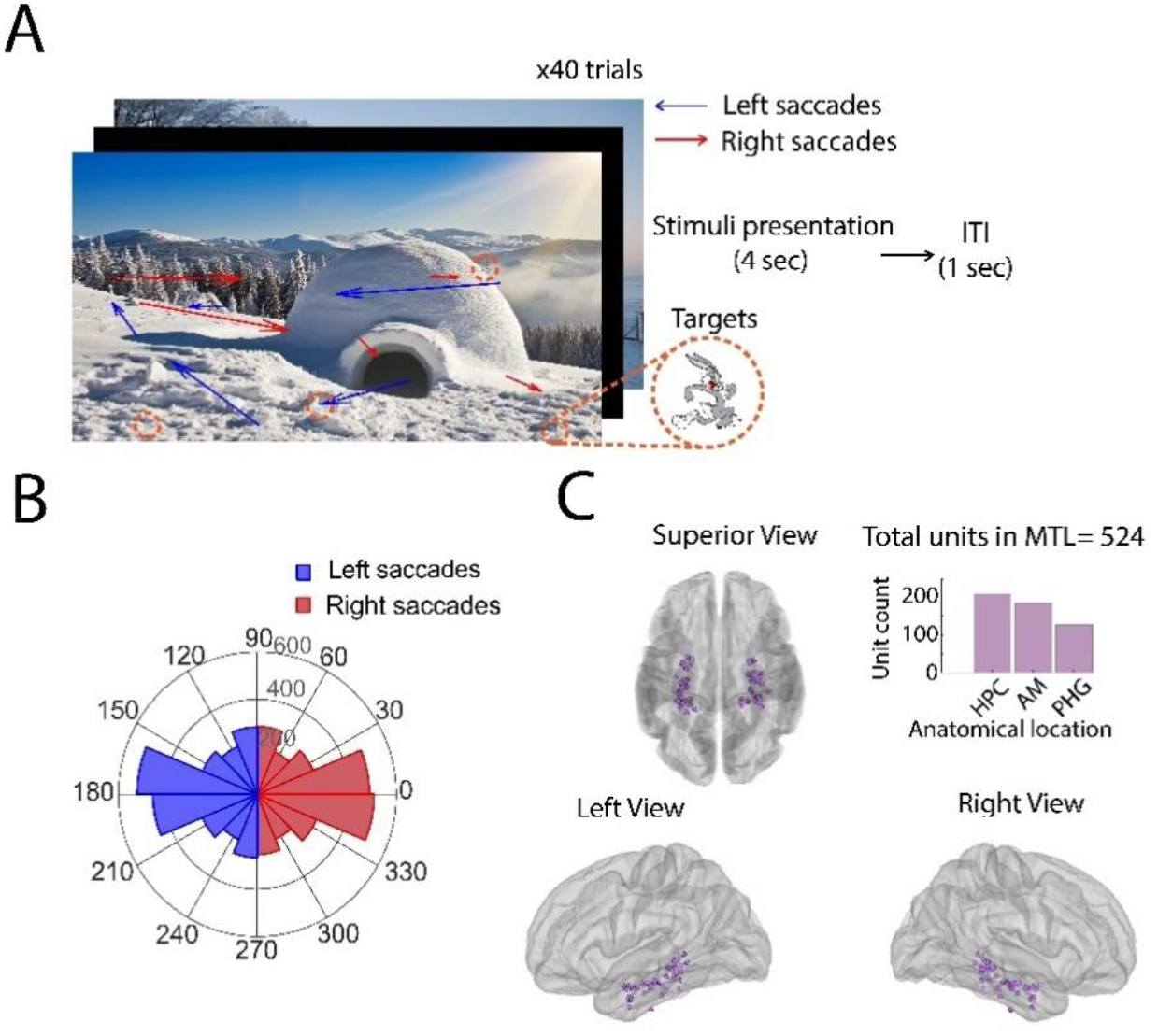
Behavioral task and recording locations. **A)** Visual search task: Participants searched for one of two target images (i.e., Bugs Bunny or Wile E. Coyote) in a series of pictures with natural scenes separated by inter-trial interval (ITI). Example of saccade trajectories (arrows) with saccades classified as leftward (blue) or rightward (red) during 4 seconds of image presentation. **B)** Leftward and rightward saccade distributions. SEM direction totals across subjects were not significantly different (paired t-test, p=0.63). **C)** Location of microwire bundles in different MTL areas. Insert: Unit counts by HPC: Hippocampus, n=210; AM: Amygdala, n=186; PHG: Parahippocampal Gyrus, n=128).

Patients reclined upright in their clinical beds with a laptop computer placed at a comfortable viewing distance in front of them. An infrared, video-based eye-tracker (EyeLink Duo; SR Research Ltd., Osgoode, Canada) was used to monitor and record their eye movements throughout the task (see below for details on eye-tracking).

### Electrophysiology

#### Acquisition and Localization

Each commercially–available, Behnke-Fried Macro-Micro electrode (Ad-Tech Medical, Racine, MN) had 8 or 9 macro electrodes along the electrode shaft and a bundle of 8 embedded microwires plus one ground/reference microwire protruding from the tip [38]. Only patients with MTL electrodes and electrodes in other anatomical locations specified by the pre-implantation hypothesis were included in this study. Data were recorded with a 128-channel Neuralynx Atlas Data Acquisition System (Neuralynx Inc, Bozeman, MT). The microelectrodes were sampled at 32kHz, with a 16-bit resolution and were bandpass filtered in hardware between 0.1 and 8kHz. Each microwire signal was re-referenced locally to one of the eight wires on the same bundle as we have done previously [39]. Electrode localization was performed by co-registering pre-op MRI with post-op CT using the iELVIS toolbox [40]. Electrodes were localized and visualized in the MTL structures across patients (Figure 1C; For occipital localizations, see Supplementary Figure S1A).

#### Spike Detection and Sorting

For offline spike detection, all microelectrode channels were bandpass filtered between 300-3000Hz. Spikes were subsequently detected using local energy measurement threshold crossings, calculated by convolving the raw signal with a kernel with an approximate width of an action potential [41]. All detected spikes were sorted using the open-source, semiautomatic template-matching algorithm OSort[38]. Similar to our previous work [39], [41], we classified clusters as putative single neurons using the following criteria (1) minimal/no violation of refractory period, (2) shape of the inter-spike interval distribution, (3) shape of the waveform, (4) separation from other clusters; and, (5) stability of firing rate (assessed by comparing the average firing rate of the neuron to multiple null distributions obtained by randomly sampling 400ms and 5s epochs from each neuron’s spike train). Groups that appeared similar to one another were merged. For extra quality control, neurons with firing rates lower than 0.07Hz were subsequently removed. Clusters that were either contaminated with noise or failed to meet the criterion described above were rejected. For the accepted clusters, the individual waveforms, along with the timestamp of each spike and its cluster definition, were saved. The insert of Figure 1C shows the total cells from each location detected in MTL (SUA from occipital locations is shown in Supplementary Figure S1A).

#### Waveform Analysis

The shape of a waveform suggests which type of neuron produced it: pyramidal neurons have more broad waveforms, and inhibitory have more narrow waveforms[42]. We followed a similar approach to Fu and colleagues to determine the classification type of spiking activity [43] from our single-unit recordings. Each neuron’s average waveform was extracted by averaging all waveforms in a specific cluster. The first trough (negative peak) and first peak times were extracted. Neurons with a trough-to-peak time of <0.5ms were considered ‘narrow-spiking’ (NS), and >=0.5ms were considered ‘broad-spiking’ (BS).

### Analysis of eye-tracking data

The SR Research eye-tracking system was used to track participants’ eye movements. The eye-tracker was placed at the bottom of the screen and connected via ethernet to a separate laptop running the behavioural task using Presentation software (NeuroBehavioral Systems, Albany, CA, USA). All participants first underwent a 13-point calibration and 9-point validation test (EyeLink Portable Duo sampled at 1000Hz, from SR Research).

Eye movements were detected using SR Research’s built-in software that uses an Identification by Velocity-Threshold (IVT) algorithm. This algorithm is a velocity-based method that separates fixations and saccades based on the distance between the current and next points [44]. To be considered a saccade, the velocity between points must exceed 30 deg/sec. When ten eye-tracking algorithms’ accuracy at detecting saccades on a sample-by-sample basis (as we do here) was compared, the IVT algorithm was second closest to human detection for saccade events [45]. An example of saccades and an image is presented in Figure 1A.

The saccade’s horizontal direction was determined by subtracting its ending point from the starting point. We coded the top left of an image as the origin, so if the saccade’s ending point was less than the starting point (a negative value), it was considered leftward. In contrast, if it was positive, it was considered rightward (Figure 1B).

### Analysis of electrophysiological data

#### Analyzing Saccade Onset Responses

Spike timestamps from offline sorted neurons were used to create binary spike trains. We then extracted 2-second epochs centred on each saccade onset for each of the sorted neurons. The spike trains were then binned into 50ms bins across trials (saccade events) to extract the instantaneous firing rate. Averages of the perisaccadic intervals (consisting of spike trains surrounding 400ms of saccade onset) were used to assess modulation (Figure 2A).

**Figure 2.**
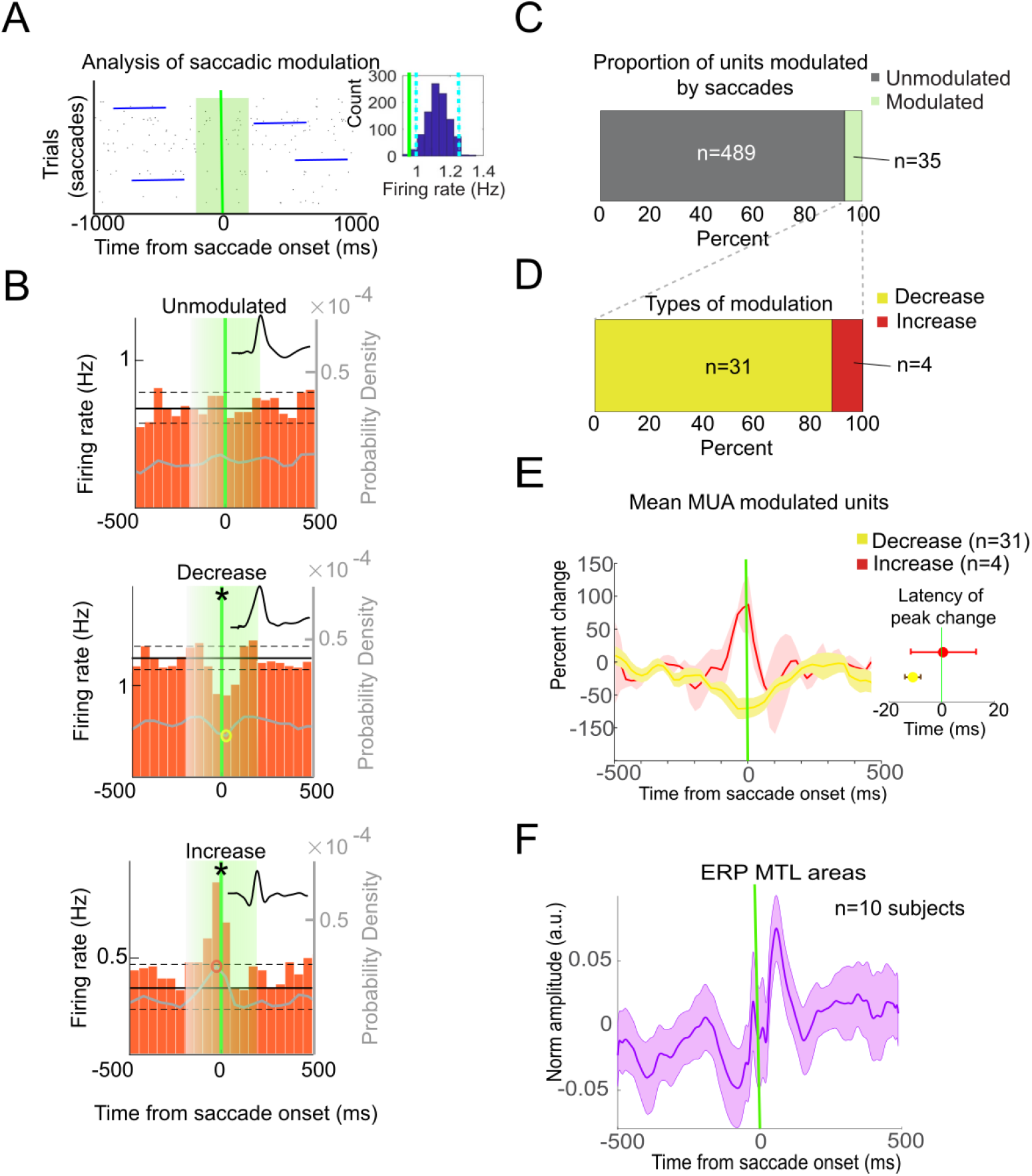
Categories and distribution of modulated units: ***A)*** The relationship between SEMs and SUA was investigated by comparing spike counts during the 400ms perisaccadic interval (green box) centred on saccade onset (green line) to those in **control periods**— bootstrapped aleatory 400ms epochs from each specific unit (dark blue lines). Insert: Null distribution of an example unit (dark blue). The green line represents the mean firing rate during the saccade period of the example neuron and the cyan dotted lines 95% confidence intervals from the random time windows. The mean firing rate (green line) is left of the lower confidence limit (cyan line), demonstrating a significant average decrease in firing rate during the perisaccadic interval. Therefore, the unit is classified as “modulated”; ***B)*** Example of the different types of modulation observed. Modulated units were subdivided into those that: i) decreased their firing rate and ii) increased their firing rate. Firing rate histograms of exemplary units and mean waveform of each unit. Yellow circle: decrease trough; Red: increase peak. Gray superimposed line indicates probability density (right y-axis). Only units with statistically significant firings during the perisaccadic periods (green box) were deemed to be modulated (asterisk); ***C)*** Distribution of units modulated by saccades; ***D)*** Distribution of the types of modulation observed; ***E)*** Mean MUA from modulated units in the MTL. Insert shows the latency of the peak change in firing rate from the MUA traces. There were no significant differences between latencies of units demonstrating an increase or decrease of firing; ***F)*** Mean ERP of electrodes in the MTL aligned to saccade onset.

To determine the statistical significance of the average peri-saccadic firing rates, we used a non-parametric permutation method. More specifically, for each session, we extracted the same number of 400ms intervals (four example intervals in dark blue in Figure 2A) as the number of saccades in the session, only at random times. By repeating this procedure 1000 times, we create a null distribution of average firing rates drawn from the same window length as the perisaccadic window and the same number of events. These eight 50ms average firing rates were averaged to create a mean perisaccadic firing rate. If the actual mean perisaccadic firing rate for a given neuron exceeded the 97.5^th^ percentile of the control null distributions, it was considered a significant perisaccadic increase. Similarly, if the actual mean perisaccadic firing rate for a given neuron was lower than the 2.5^th^ percentile of the control null distribution, it was considered a significant perisaccadic decrease (Figure 2A inset).

As a result, neurons were classified into one of three groups – (1) non-modulated (no significant increase or decrease in firing rate during the perisaccadic interval), (2) perisaccadic increase (significant increase in firing rate during the perisaccadic interval), and (3) perisaccadic decrease (significant decrease in firing rate during the perisaccadic interval (Figure 2C). The multiunit activity (MUA) was calculated by averaging all modulated neurons’ average firing rate aligned to saccade onset, separately for each category (decrease or increase). The percent change was calculated based on the ratio of the average firing rate per neuron during the saccade period, divided by the mean firing rate in the randomized control periods (Figure 2E). Then that percent change was averaged across all neurons. We also calculated the perisaccadic firing probability density by fitting all spike timestamps in 2 seconds around each saccade with a kernel distribution with a bandwidth of 50ms. This probability density was then upsampled (using a bicubic spline interpolation) to obtain a 1ms resolution. The maxima and minima of this upsampled firing probability density were used to calculate the peak and trough latency of the firing rate increases and decreases, respectively, for the saccade-modulated units (Figure 2E insert). A histogram of the latencies is also presented in Supplementary Figure 2C. Subsequent statistical tests on distributions based on these classifications were determined to be parametric (paired t-test, one sample t-tests) or non-parametric (Wilcoxon Signed-Rank Test or Wilcoxon Rank-Sum Test), based on the results of the Lilliefors normality test.

Although the interest in the present study focused on SU saccadic modulation of MTL structures, in two participants, we had the unique opportunity to simultaneously record activity from the occipital cortices (OCC). Therefore, we performed a unit-by-unit analysis from neurons acquired in this area, shown in the supplementary material, in a similar way as MTL neurons. A fourth classification was included for OCC: rebound modulation, consisting of an observed initial decrease followed by a sharp increase. We calculated the minima and maxima of the average firing rates in the 400ms perisaccadic window and compared these to randomly generated control null distributions of maximum and minimum average firing rates. If the maximum perisaccadic firing rate for a given neuron exceeded the 97.5th percentile of the control null distributions, and the minimum perisaccadic firing rate was lower than the 2.5th percentile of the control null distribution, it was considered a rebound (Figure S2C). No units from the MTL presented a characteristic rebound.

To determine if the amplitude of the post-saccade event-related potentials (ERPs) [15] was related to firing rate changes, post-saccade ERPs were computed from voltage traces of the most distal macroelectrode (i.e., the one closest to the corresponding microwires). Each macro electrode had a 200Hz lowpass filter and was downsampled to 1000Hz. A 60 Hz notch filter was applied to reduce environmental noise from the signal. Each macroelectrode ERP was normalized by dividing by the standard deviation of the voltage trace of that electrode during the entire recording. The ERPs found to have a reversed polarity were flipped to obtain a positive waveform (MTL= 1/58; OCC= 5/8). We eliminated responses from electrodes with large artifacts in the ERP and, therefore, considered open or broken channels (MTL= 4/58; OCC= 0). The subsequent average from all electrodes was calculated and plotted for MTL (Figure 2F) and occipital electrodes (Supplementary Figure S1G). The root mean squared (RMS) amplitude of the normalized ERP was then calculated from 10ms-510ms post saccade onset. This time was chosen as it is in line with the significant portions of the saccade ERP from our earlier work [15].

Previous LFP work has identified differences in responses based on the direction of saccade relative to recording location [30], even as it relates to potential oculomotor artifact [46]. Therefore, we further labelled saccades as ipsiversive if the saccade’s direction was towards the side of the recording electrode’s hemisphere or contraversive if the saccade’s direction towards the opposite hemisphere. We repeated the categorization and ERP analyses described above, now separating events based on the saccade direction. Additionally, a paired t-test was used to compare ipsiversive to contraversive saccade RMS distributions. Finally, separating ipsiversive and contraversive saccades, we calculated the Spearman rank correlation between the ERP RMS amplitude and the percent change in firing (decreases only) of the corresponding modulated neurons close to those electrodes. Single unit modulation was similar across the MTL structures, so they were grouped together for further analysis (Supplementary Figure 2).

#### Analyzing Image Onset Responses

Single unit responses to image onset were analyzed in a window between 200 and 1700ms after image onset, in line with previous single-unit literature [47]. Neurons were marked as modulated by image onset if their mean firing rate in this window significantly increased or decreased compared to the corresponding null distribution (Figure 4A). Control null distributions for image onset analysis consisted of 1500 ms periods of spike trains at randomized timepoints. The MUA was also calculated for neurons that increased or decreased their firing rate to image onset (Supplementary Figure 3).

## Results

### SUA within the MTL are modulated in the peri-saccadic interval

We recorded 524 neurons from MTL regions (hippocampus, amygdala and parahippocampal gyrus) across 11 patients. Table 1 shows the distribution of neurons analyzed from the different areas. Waveforms of some example neurons are shown in Figure 2C. In two participants, we had the unique opportunity to simultaneously record activity from the occipital lobe (Table 1, Figure S1C). Overall, the various MTL structures presented similar results and therefore were grouped (Figure S2). To determine whether saccades modulated single-unit activity in the MTL on a unit-by-unit basis, we analyzed the firing rate changes in a 400ms window surrounding each saccade and compared this activity to a matching null distribution generated by selecting random windows throughout the recording session (see Methods for details). In doing so, we found that 6.7% (35/524 neurons) of MTL neurons were strongly modulated by the saccade (Figure 2B). This percentage is somewhat lower (~28%) than in another study investigating eye movements in humans [20]. However, factors such as saccade detection and modulation criteria differ between these studies [20]. When separating by direction (See Saccadic Modulation of the MTL varies with saccade direction), the proportion of modulated neurons found is in line with previous work with primates (20%) [28]. Interestingly, a more significant proportion of occipital lobe neurons were modulated around the time of saccades (46.4%, 58/125 neurons; Figure S1B).

### SEM related modulation is predominantly inhibitory

To determine how MTL SUA is modulated in the peri-saccade interval, we divided each unit into one of three categories: unmodulated, firing rate increase, or firing rate decrease. Of the saccade modulated units in the MTL, we found that most units demonstrated a reduction in firing rate within the perisaccadic window (88.6% n=31/35), with the small remainder demonstrating increased firing rate (11.4%; n=4; Figure 2D).

To further interrogate this perisaccadic modulation, we classified the units into two groups based on their waveform width: broad-spiking (putative pyramidal cells) and narrow-spiking (putative interneurons). We found that of the small population of neurons that did increase their firing rates in the peri-saccadic window, a majority were the putative interneuron group of narrow-spiking units (75%, 3/4) (Figure 3). In contrast, among the more prevalent group with decreased firing rates, most neurons were broad-spiking (77.4%, n=24/31). The overall distribution of broad and narrow-spiking units for MTL can be seen in Figure 3. These findings suggest that saccades predominantly decrease the activity of putatively excitatory neurons and predominantly increase the activity of putatively inhibitory neurons.

**Figure 3.**
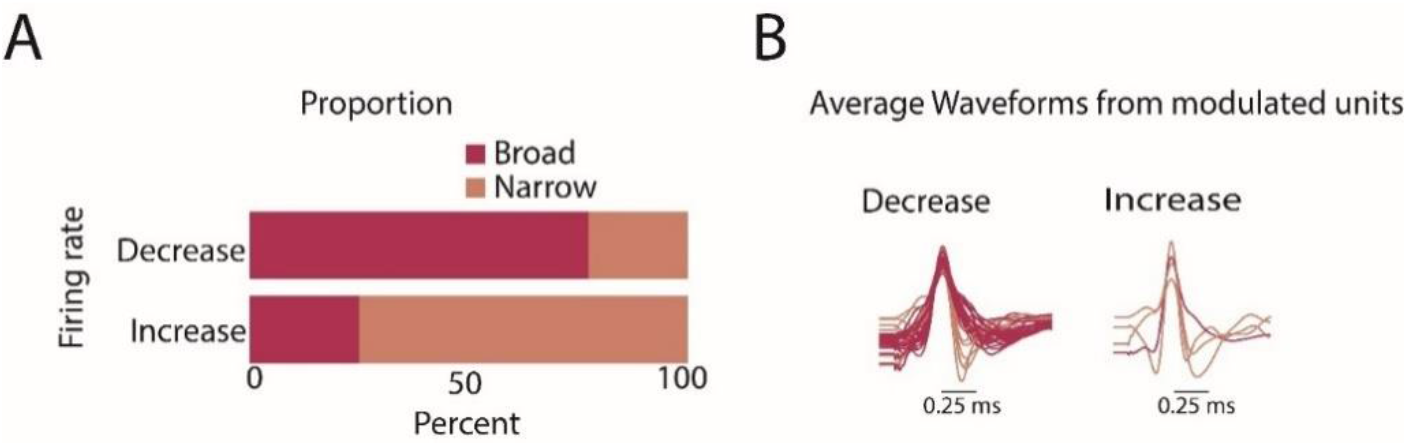
Broad spiking units largely demonstrate decreases in firing rate. **A)** Proportion of ‘broad’ and ‘narrow’ spiking separated by the type of firing rate changes observed in the perisaccadic interval. **B)** Average normalized waveforms from the MTL areas separated by type of firing rate change.

In contrast, many neurons in occipital regions demonstrated a general decrease in firing rate, like the modulated MTL neurons (43.64%; n=24/55). 5 larger fraction demonstrated a characteristic rebound activation pattern in the perisaccadic window (56.36%; n=31/55), which consisted of a significant transient decrease in firing in the peri-saccadic window, followed by a sharp, significant increase (Supplementary Figure S1C). Only a small subset of the modulated occipital lobe neurons demonstrated an increase in firing rate (5.45%, n=3/55) (Supplementary Figure S1D). Akin to the modulated units in MTL structures, units in the occipital lobe that demonstrated firing rate increases were mostly narrow-spiking (66.7%, n=2/3). Interestingly, most rebound-modulated neurons in the occipital lobe were narrow-spiking (80.64%, n=25/31; Supplementary Figure S1F). As with the MTL, the majority of occipital lobe neurons that decreased their firing rate were broad spiking (87.5%, n = 21/24).

### Saccadic modulation precedes SEM onset

Our previous finding that saccade-related ERPs are aligned to saccade onset rather than offset [15], as well as the contemporary understanding of the arrival time of CDs relative to saccade onset [31], together suggest SUA should be modulated before and/or during a saccade [31]. To investigate this, we separated the modulated units in the MTL by their modulation type (increase or decrease). We obtained the maximum average firing rate latency in a 400ms window surrounding saccade onset for units that up, modulated. Similarly, we found the latency of the minimum of the average firing rate in the same window for the units that were down-modulated. In doing so, we found that the trough of the firing rate decreases occurred at −11±3ms relative to saccade onset, whereas the peak of the firing rate increases occurred at 0.5±12.57ms relative to saccade onset (Figure 2E inset). Only down-modulated neurons were significantly different from zero (up-modulated, t-test p=0.97; down-modulated sign rank p=0.001) though these latency distributions were not different from each other (rank-sum, p=0.33). In summary, that the perisaccadic modulation occurs before or during saccade initiation, well before the hippocampus would be expected to respond to changes in visual input, is consistent with a CD-like modulation of MTL units [34].

Similarly, firing rate troughs and peaks in the occipital lobe occurred at −11ms ±3.2ms and 0.33ms ± 15.7ms relative to saccade onset. However, for units that demonstrated a rebound pattern, the rebound increase occurred at a longer latency (27 ± 3ms; Figure S1E). Thus, SEM-associated SUA modulation in both the MTL and occipital regions occur similarly before or during SEM, unlike image onset modulations of SUA that occur much later (see below and also [34]). This pattern is consistent with the notion that this modulation of SUA reflects the influence of a CD-like signal [48].

### The units and the response timing of the SEM-related SUA are distinct from those from image onset

The distinct nature of the ERPs associated with SEMs and image onsets [15] implies that the SUA associated with SEMs and image onset should also be separable. Specifically, ERPs associated with image onset were of an evoked nature, suggesting that we should observe increases in SUA instead of the inhibition-dominated changes in SUA associated with SEMs. Furthermore, the image onset evoked ERP’s temporal evolution was slower, and thus we would expect firing rate changes to occur later.

To explore these possibilities, we analyzed peri-image onset responses in a window of time between 200ms and 1700ms after image onset [47]. Neurons were classified as being modulated by image onset if their firing rate in this window significantly increased or decreased compared to the corresponding null distribution (Figure 4A). We found that 13.7% of the recorded MTL units were modulated by image onset (n = 72/524; 67/524 increase; 5/524 decrease) (Figure 4B). This fraction is similar to that reported in previous studies [33]–[36], [39].

**Figure 4.**
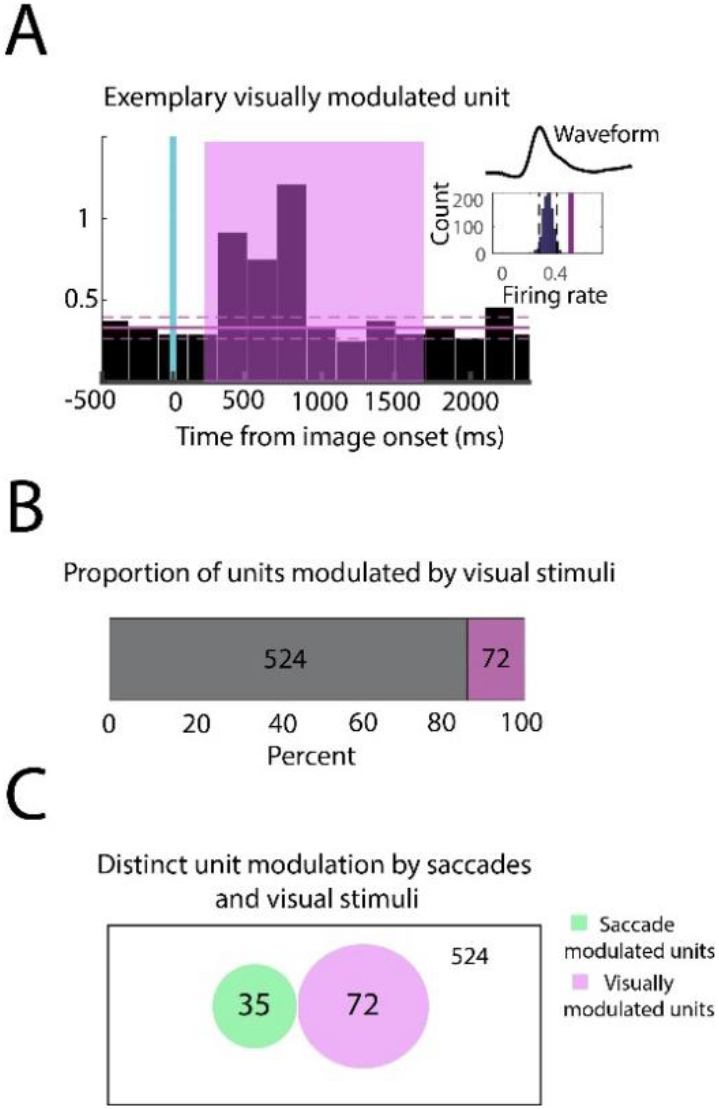
MTL units modulated following image onset are distinct from those modulated during SEMs: **A)** Histogram of an example unit modulated by image onset noted by the increase in firing rate starting 500ms after image onset (blue line). Insert shows the waveform and mean firing rate during the analysis period (200-1700ms); **B)** Proportion of units modulated (purple) by image onset; **C)** Venn diagram of visually modulated units (purple) or perisaccadic period (green). No intersection was found between the two populations of units, suggesting that units modulated by saccades are distinct from those visually modulated.

Interestingly, we found the population of neurons modulated by image-onset to be distinct from those modulated around the time of SEMs (Figure 4C). Furthermore, image-onset, modulated neurons demonstrated a peak increase in firing rate at 650 ± 100ms after image onset, well after the saccade-related units (Figure S3). Thus, within the MTL, the units receiving visual information following image-onset appear distinct from those responding to saccade-related information. A higher percentage of units in the occipital cortex than in the MTL were modulated following image onset (17.6%; n=22 /125) (Figure S1H). Unlike the MTL, 4% (5/125) of occipital units were significantly modulated both following image presentation and during SEMs.

Taken together, these results demonstrate a clear dichotomy between SEM-related and image-onset unit activity in the MTL, where the former is dominated by inhibition, begins and ends earlier, and is mediated by a signal that modulates a unique set of neurons. The latter feature of the SEM-related units in the MTL suggests that the modulating signals associated with SEMs within the MTL target distinct cell populations than units receiving visual input (Figure 4C).

### Saccadic Modulation of the MTL varies with saccade direction

In other SEM-related CD circuits, a CD signal conveys information regarding the ensuing saccade’s direction and magnitude, likely facilitating anticipatory remapping of receptive fields in visual areas [49], [50]. Thus, we would expect that if a CD signal mediates the MTL firing rate modulation, we should observe directionally selective modulation in the peri-saccade interval. To investigate this question, we separated saccades into two groups, ipsiversive and contraversive (there were no significant differences between the number of saccades in these groups paired t-test p=0.16; Figure 1B). In addition to the 6.7% (n=35/524) of neurons that were classified as modulated when collapsing across saccade direction (Figure 2B), separating saccades into ipsi- and contraversive groups revealed a much larger fraction of modulated neurons (20.4%, n=107/524) (Figure 5B). In fact, most modulated units when analyzing all saccades (28/35) were also directionally modulated (i.e., modulated in one particular direction), suggesting that the previous categorization of SEM associated SUA modulation was masked by pooling together all the saccades. This finding provides strong evidence that saccadic modulation of neural activity in the medial temporal lobe structures is directionally dependent.

**Figure 5.**
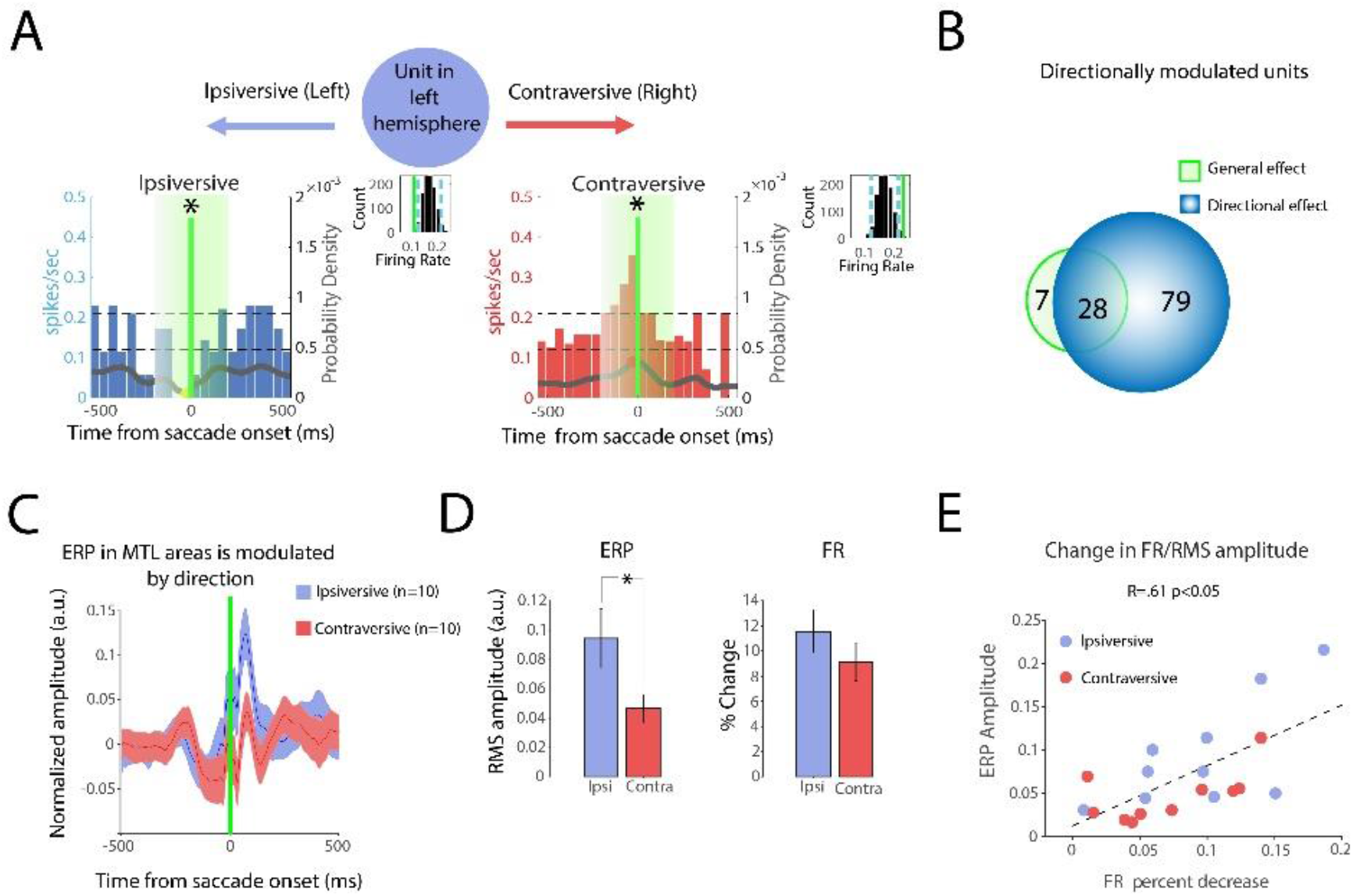
SEM-related SUA and ERPS are directionally modulated and correlated to one another. **A)** Ipsi- and contraversive firing rate changes of an example unit. Units were classified as “modulated by direction” if there was a significant increase or decrease associated with contraversive or ipsiversive saccades. The unit in this example shows a decrease in ipsiversive saccades and an increase in contraversive saccades. Insert shows the distribution of units during control periods and mean firing rate during the saccadic period (green line). **B)** Venn diagram showing the proportion of units modulated by saccades (green) and the units that showed a directional effect (blue). **C)** ERP from electrodes localized in the MTL associated with ipsi- and contraversive saccades. **D)** The left graph shows LFP RMS amplitude separated by the direction of the saccade. RMS amplitudes of ipsi- and contraversive ERPs are statistically significantly different. (Paired t-test, *p<0.05). The right graph shows the percentage modulation of units during the saccade period compared to randomized control periods. **E)** Correlation of LFP and unitary activity measurements sowed in A, presenting a positive correlation.

To further characterize this directional modulation, we classified these effects as 1) Lateralized decreases (a decrease in firing rate, but only for ipsiversive OR contraversive saccades); 2) Lateralized increases (an increase in firing rate, but only for ipsiversive OR contraversive saccades); 3) Mixed (increased firing rate in one direction, and decreased firing rate in the opposite direction). In the MTL, most directionally modulated units fell into the lateralized decreases category (n=73/107), consistent with one of our main findings, that units are largely inhibited in the peri-saccadic interval. A smaller subset of units had mixed modulation (Figure 5B; n= 23/107). Only a small population of neurons presented lateralized increases (n= 11/107) (Supplemental Figure 4A).

As before, our cell-type analysis revealed that the majority of MTL neurons that presented lateralized decreases in firing were excitatory broad-spiking (70%; n=51/73), while a majority of the neurons that presented a lateralized increase in firing were inhibitory narrow-spiking (64% n=6/11). The majority of units presenting mixed directional effects were classified as “broad-spiking” (65.21%; n=15/23) (Supplemental Figure 4B).

Additionally, we investigated the directional modulation of units in the occipital lobe. Here, 62/125 (49.6%) units were directionally modulated. Unlike the MTL, there was a smaller overlap between the directionally-modulated units and those units modulated when collapsing across saccade direction (n = 25 units were common between both groups) (Figure S1I). Overall, there was significant directional modulation of single-unit activity in the occipital lobe, similar to that observed in the MTL.

### SEM related SU firing rate decreases correlate to post-saccade ERP amplitude

Our previous work identified a short-latency, saccade-aligned phase-resetting ERP in the MTL LFP aligned to the saccade onset [15]. The combined observation that post-saccade ERPs align to saccade onset [15] and that SU firing rate changes are confined to the peri-saccade interval suggests that the SU firing rate changes drive the subsequent ERP. If this is so, then the amplitude of the SEM-related ERP should depend on the saccade’s direction and be correlated to SU firing rate changes. To explore these possibilities, we calculated the saccade-aligned ERP for both ipsiversive and contraversive saccades (Figure 5C). We found that the RMS values 10-510ms after saccade onset (corresponding to the significant portions of the saccade ERP from our previous work) depended on saccade direction (Figure 5D). Specifically, the ipsiversive saccade ERP had a significantly greater RMS magnitude than the contraversive ERP (Ipsiversive = 0.09 ±0.019 normalized amplitude; Contraversive = 0.04 ±0.009 normalized amplitude; p<0.05 Figure 5D). Thus, both the ERP and SU firing rate changes depend on the direction of the saccade.

Additionally, if the earlier peri-saccadic firing rate decreases underlie the generation of the later occurring post-saccade ERP, we would expect firing rate changes to be correlated to ERP amplitude. To explore this possible relationship, we measured firing rate decreases in the directionally-modulated MTL units (contraversive or ipsiversive modulated unit) and corresponding ERP (contraversive or ipsiversive ERP from electrode corresponding to the modulated unit) RMS amplitude for each patient. One patient was eliminated from this analysis as no modulated units were found. Spearman’s rank correlation between these measures revealed that firing rate decreases were positively correlated to ERP magnitude (Figure 5E). Thus, the more strongly BS neurons were inhibited – the cell-type that demonstrates the most consistent decrease in firing rate – the larger the ERP. We infer from this relationship that the post-saccade ERP [15] may be, in fact, a proxy for the strength of an earlier, largely inhibitory modulating signal (see Discussion).

## Discussion

In our previous work [15], we presented LFP evidence that 1) the MTL is uniquely modulated around the time of SEMs as compared to image onset; 2) this modulation was consistent with a phase resetting response, and; 3) the alignment to saccade onset of this phase resetting was suggestive of a CD-like signal. These conclusions comported well with extensive NHP literature regarding the extra-retinal contributions to the modulation of LFPs in MTL and other visual areas [13], [14], [20], [30], [48], [51], [52]. Here we present six key findings derived from human SU recordings that greatly extend these findings to provide a plausible mechanism by which extra-retinal signals accompanying SEMs modulate the hippocampus and surrounding MTL structures in humans. Our six key findings are: (1) the peri-saccade interval is dominated by inhibition (firing rate decreases of BS units, firing rate increases of NS units); (2) SU activity is modulated largely before and during the saccade; (3) the modulation contained directional information; (4) distinct population of neurons are modulated by saccades and image onset, and saccade-related modulations occur much earlier; 5) saccades modulation of occipital SUA is distinct from MTL SUA modulation; 6) the amplitude of the post-saccade ERP is correlated to the magnitude of the firing rate decrease of BS units in the peri-saccadic interval. Through the manuscript, we grouped MTL results based on similar modulation patterns (Supplementary Figure 2), suggesting a CD like signal is widely present within the MTL and thus likely generally relevant to mnemonic processes, albeit different structures having variable roles in memory[35] [53]. However, given that the majority of units we observed within the MTL were from the hippocampus and the extensive knowledge about hippocampal physiology and function, we focus our discussion specifically on hippocampal circuitry as an exemplar.

### Firing rate decreases dominate the peri-saccade period

The combination of increased NS and decreased BS firing rates suggests that the incoming saccade-related signal has a net inhibitory influence. We argue that this pattern is consistent with a general CD motif, in which motor-related signals excite local inhibitory interneurons, which then inhibit excitatory neurons [16], [17]. There are other possibilities. It could be argued that decreases in BS activity arise from the reductions in afferent visual input during saccadic suppression [25]. Arguing against this possibility, however, BS firing rates change before afferent visual input would arrive in the MTL [34], [54]. Moreover, the accompanying increases in NS unit activity would be hard to explain with decreased afferent drive. Another possibility is direct inhibition through long-range GABAergic projections. However, the direction of the SUA modulation we observe makes this unlikely [55].

Observing this saccade-related inhibitory mechanism in single units sheds light on the post-saccade ERP that we [15] and others [13], [14], [20], [51] have characterized. Since inhibition alters the timing of ongoing spiking through a pause-rebound mechanism [56], pausing MTL circuits right before a saccade would orchestrate synchronized periods of excitability following the saccade. And, because these MTL circuits are tuned to resonate at theta [57] as we have observed in the LFP [14], [15], these dynamics would align activity of a 3-8Hz oscillation that is phase-locked to the initiation of a saccade [15]. Notably, the resulting “phase-reset” need not reflect the modulation of ongoing oscillations – an important consideration because continuous oscillations like theta are not prominent in the primate hippocampus [58]–[60] and certainly not continuous like those in the rodent [61]. We also provided evidence linking SU inhibition to ERPs. Because larger inhibitory post-synaptic potentials increase the probability of consistently timed rebound spikes [56], greater saccade-related SU inhibition should drive larger ERPs. This relationship is precisely what we observed when correlating the two. Thus, by combining SU recordings with ERP analyses, we provided much needed insights into *how* saccadic eye movement could initiate the observed LFP modulation within the theta frequency range.

Beyond eye movements, such an inhibition governed hippocampal phase reset is well described in response to both alerting stimuli in rodents (auditory clicks, smell, touching) [62]–[64] and electrical stimulation of intrahippocampal pathways [65]–[67]. Considering these parallels, perhaps eye-movement plans act as an alerting signal, preparing the hippocampus for important new visual input. Indeed, hippocampal phase resetting temporally organizes LTP and LTD periods relative to the stimulus generating the reset [68]. Thus, saccades, along with a myriad of externally driven salience signals, may leverage a general mechanism – inhibition mediated phase resetting – to organize the encoding of new and important information. This link between active visual sensing and specific memory processes in mnemonic structures is consistent with a large literature rife with evidence linking saccadic eye movements to memory strength (for review, see [5], [69], [70]).

### SU activity reflects the early arrival of spatial and temporal parameters of ensuing eye movement

We further evaluated two core CD properties: that they should occur before movement and contain spatial information about the ensuing movement [31]. We indeed found that MTL SUs were predominantly modulated in the pre-saccadic window. This comports well with our previous observation that MTL LFP modulation is aligned to saccade onset and not its termination (i.e., fixation) [15]. Both findings are consistent with the notion that MTL neurons are more modulated by the motor planning of saccades than the resulting change in visual input. We also found strong directional modulation in MTL SUA. So much so, that collapsing across saccade direction drastically underestimated the proportion of saccade-modulated units (6.7% of all MTL units) compared to separately analyzing ipsiversive and contraversive movements (20.4%). Moreover, we found that ipsiversive saccades were related to larger ERPs than contraversive in the MTL, consistent with previous research in NHPs comparing temporal and occipital lobes [30]. In that study, the direction was also associated with significant differences in temporal lobe LFP but not occipital [30]. In our study, a vast majority of saccade-modulated MTL units were directionally modulated (107/114, 93.9%). Although directionally modulated units were also present in the occipital lobe, they represented a relatively smaller proportion of the total number of modulated occipital units (62/87, 71%). It has been suggested that the modulation of units directly receiving visual information reflects an exafference signal more than the CD signal [71]. Accordingly, our MTL findings may reflect the outcome of a more unadulterated CD, whereas the occipital lobe saccade modulation is likely the summed influence of a CD and visual reafference [30].

### SU modulated by saccades are distinct from SU modulated by image onset

Visually responsive neurons increased their firing rates long after saccade modulated neurons, with their MUA peaking ~500ms after image presentation. SU also exclusively increased their response to image onset, consistent with an evoked response [15] and sharply contrasting with the decreases in SUA associated with SEMs. Surprisingly, image onset and saccades modulated separate sets of MTL neurons, suggesting that the pathways mediating the effects, and their targets, are distinct. In the neocortex, modulating influences exert themselves in extra-granular layers while driving thalamic inputs exert their influences primarily in the granular layers [72]. Though it is hard to translate neocortical architectonics to the hippocampus, we conject that image onset responses reflect entorhinal inputs that excite neurons in the pyramidal layers [73]. In contrast, elucidating the source of the modulating CD signal, which excites interneurons, will require additional experiments.

For CD modulation to originate from motor structures (i.e., the superior colliculus), there must be anatomical connections between the oculomotor systems and the MTL. While there are no direct anatomical connections between these systems, indirect cortico-cortical connections may facilitate this type of communication. For instance, there are indirect connections between frontal eye fields (FEF) and the hippocampus [9]. This network supports top-down attentional control and is partly responsible for generating voluntary eye movements. Other voluntary controllers of eye movements, such as the superior colliculus [74] and the pulvinar, may indirectly influence MTL structures[75]. Another possible source of both directional information and inhibition within the hippocampus is the nucleus reuniens which contains head direction neurons in rodents [76], appears to have a modulatory influence within the CA1 region mediated predominantly through inhibition [77], [78], and is a critical memory related node mediating communication between the hippocampus and the medial prefrontal cortex [79]. Recently, saccade direction cells have been demonstrated in the NHP entorhinal cortex [28] – another target for reuniens efferents [80].

### Role of directional information in the MTL

How might a CD signal be utilized in the MTL? Rather than indiscriminately inhibiting BS units, our results suggest that the CD could inhibit (or excite) distinct MTL units depending on the associated saccade’s direction. This spatial tuning is reminiscent of the spatial and directional information that MTL structures in rodents use to create maps of the physical world, or cognitive maps more generally [81], [82]; place cells [81], grid cells [83], [84], and head direction cells [85] have been demonstrated in the MTL. Primates, by contrast, primarily explore their physical world with their eyes. It is, thus, unsurprising that spatial information about eye movements should be reflected in the primate MTL SUA [14], [18], [28], [29].

Here, we add to this literature by showing that this directionally tuned peri-saccadic modulation is primarily inhibitory, suggesting that its role may be akin to the well-accepted role of CDs in stabilizing visual perception [21], [27], [86]. Specifically, CDs are thought to “unite separate retinal images into a stable visual scene,” as was beautifully demonstrated in experiments that inactivate the CD while NHP explore visual scenes [27]. The resulting alterations of visual perception provided early links between CD and perception. The neuronal mechanism underlying such a perceptual role for a CD, was speculated to be akin to the anticipatory shifting of receptive fields in the lateral interparietal sulcus during saccadic eye movements [49]. Analogously, the directional information (i.e., the “path” in path integration) we identify in the human hippocampus could be used to stitch together sequentially sampled windows into a singular memory of a scene[11], [12], [87]. Perhaps this mechanism may even generalize to reflect the hippocampus’ emerging role in encoding “conceptual maps” [82], [88]. Whether initiated by the traversal of visual or conceptual space, a targeted inhibitory-rebound mechanism could facilitate a construction from parts, reminiscent of episodic memory semanticization [88], through path integration.

### Limitations of current work

If the peri-saccade modulation is solely a result of extra-retinal signals, it should persist in darkness. While the restrictions of our clinical setting precluded our test of this hypothesis, previous work with human and NHP suggests that both saccade-related ERPs and SUA modulation are observed in darkness and/or blank screens [18], [20], [19], [51] but see [14]. For example, in humans, rapid-eye movements in sleep and wakefulness generated both ERPs and SUA modulations in the perisaccadic period. The pattern was also characteristic of decreases in SU firing rates before the rapid eye movement with increases afterwards in all cases other than awake non-visual stimulation[20]. Their observed rebound pattern is reminiscent of the firing rate change observed exclusively in occipital lobe units (Supplementary Figure 1), a result that might have arisen from averaging of up and down-modulated units.

By contrast, a recent NHP study demonstrated no appreciable modulation in hippocampal MUA following saccades on a blank background, despite saccade-associated phase clustering in the LFP [51]. However, this analysis approach averaged across all units, rather than focusing on the small proportion of modulated units, as we and others have done. Had this more sensitive approach been used, SUA modulation may have been observed while looking at a blank screen. Relatedly, our initial investigations only revealed phase clustering when exploring images, not during saccades on a dark screen [14]. However, this lack of clustering was based on saccades made during a blank screen in the inter-trial interval immediately following reward delivery, rather than longer periods of complete darkness outside the task regime. Also only LFP data was analyzed without SU activity, and the analysis was from data collected from a single NHP. Thus, while there is some heterogeneity in the literature, a large body of work supports the hypothesis that saccadic modulation of activity in the MTL persists in the absence of visual input, further supporting the notion that this type of modulation arises from an extra-retinal signal.

## Conclusion and outstanding questions

In summary, we characterize SUA in MTL and occipital lobe structures to explore the hypothesis that an SEM-related CD-like signal exists in mnemonic structures of the human brain. Employing a unit-by-unit analysis of firing rate changes, we provide compelling evidence that SEMs are associated with a modulating influence characterized largely by inhibition and containing directional information. We show that saccade related modulation of MTL units occurs earlier than image onset modulation and is more inhibitory and directionally-tuned in the MTL than in the occipital lobe units. All these distinctions are consistent with MTL neurons receiving a saccade-related CD signal.

We hope that these discoveries inspire future work to investigate how a CD-like signal contributes to memory, ideally by interrupting the CD signal [27]. The next steps in this pursuit would involve elucidating the pathway by which CD-like signals reach MTL structures, and understanding the inhibitory influence of this pathway. Beyond clarifying the circuitry mediating hippocampal saccadic modulation, cell-type and circuit explorations could inform the exciting field of neuromodulation. To date, such studies have been limited to loss-of-function results following electrical stimulation of hippocampal structures (i.e., memory disruption [89]–[91]). A deeper understanding of how hippocampal circuitry is recruited during visual search may provide new avenues for cell-type and timing-specific approaches to neuroprosthetic devices for memory.

## Author Contributions

Conceptualization, C.K., and T.A.V.; Methodology, K.P., C.K., A.G.P.S; Software, C.K., K.P., and A.G.P.S.; Formal Analysis, A.G.P.S, K.P. and C.K.; Investigation, C.K. A.G.P.S and K.P.; Data Collection, C.K. A.G.P.S, K.P. and V.B; Single Unit Selection, A.G.P.S Visualization, A.G.P.S, K.P. and C.K.; Writing – Original Draft, C.K., A.G.P.S and K.P.; Writing – Reviewing and Editing, C.K., A.G.P.S, K.P., K.H., K.D. and T.A.V.; Surgical Implantation, S.K and T.A.V.; Funding Acquisition, T.A.V.; Resources, T.A.V.; Supervision, T.A.V.;

## Acknowledgements

We would like to acknowledge Azadeh Naderian and Harish Babu who helped with some of the initial sorting of the single units.

## SUPPLEMENTARY FIGURES

**Supplementary Figure 1.**
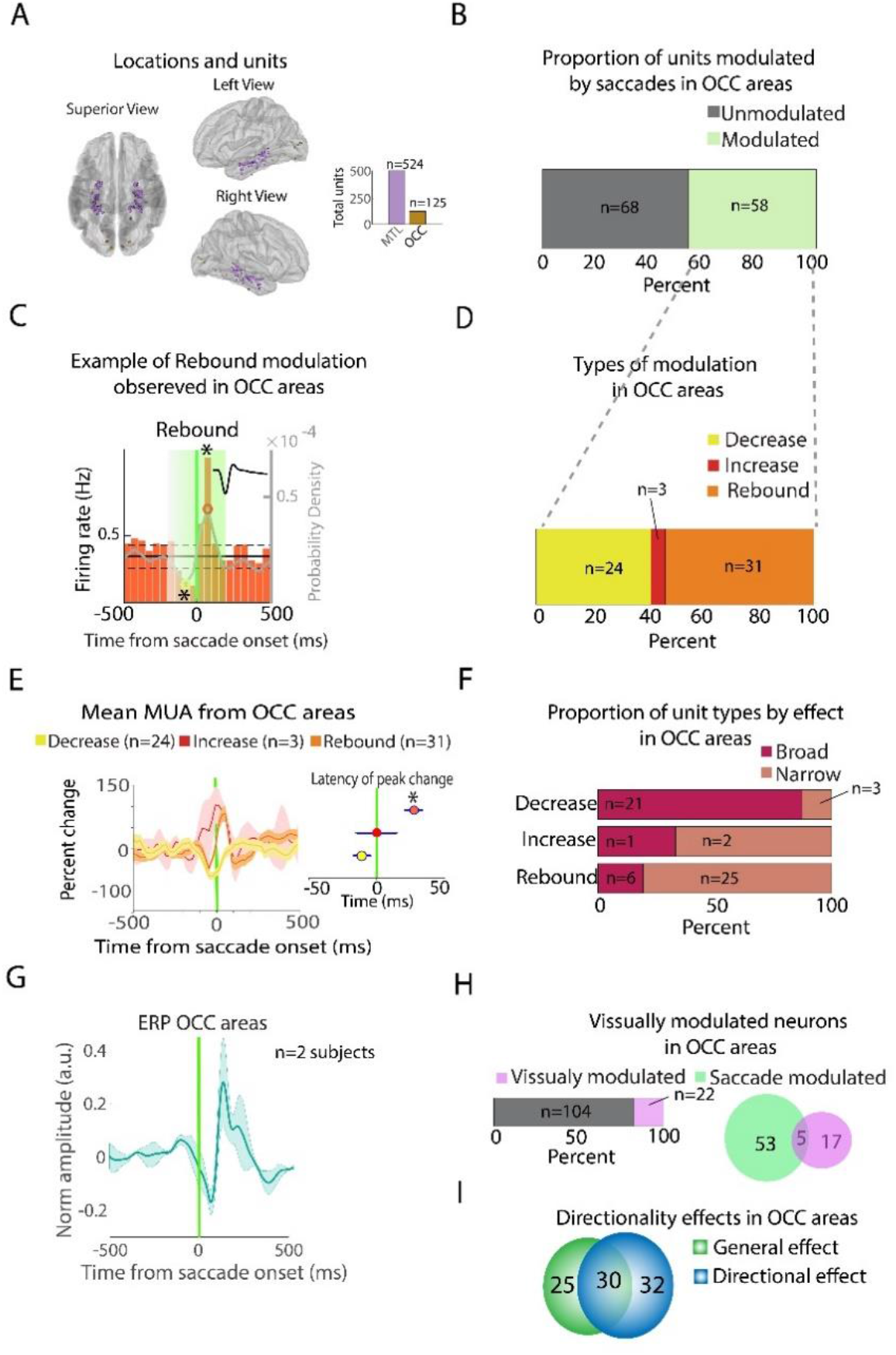
Modulatory effects in Occipital areas. **A)** Electrode locations in occipital areas from 2 patients. Insert: Unit count in the OCC areas compared to the MTL areas; **B)** Distribution of units modulated by saccades; **C)** Firing rate histogram of exemplary unit presenting a Rebound Type of modulation observed only in OCC areas (mean waveform of the unit in top right insert). Asterisks represent significant decreases (yellow circle) or increases (red circle) within the perisaccadic period (green box). Gray superimposed line indicates probability density (right y-axis) for visualization purposes. **D)** Distribution of the different types of modulation observed as explained in the methods section. **E)** Mean MUA from modulated units in the OCC. Insert shows the latency of the peak change in firing rate from the MUA traces; **F)** Proportion of ‘broad’ and ‘narrow’ spiking separated by the effect in firing rate in the perisaccadic interval; **G)** Mean ERP of electrodes in the MTL aligned to saccade onset; **H)** Left: the proportion of visually modulated neurons in the OCC; Right: Venn diagrams show the proportion of units visually or saccade modulated. Note that there is a significant overlap of units responding to both events in the OCC; **I)** The majority of saccade modulated neurons also present a directional modulation. MTL, Medial Temporal Lobe; OCC, Occipital.

**Supplementary Figure 2.**
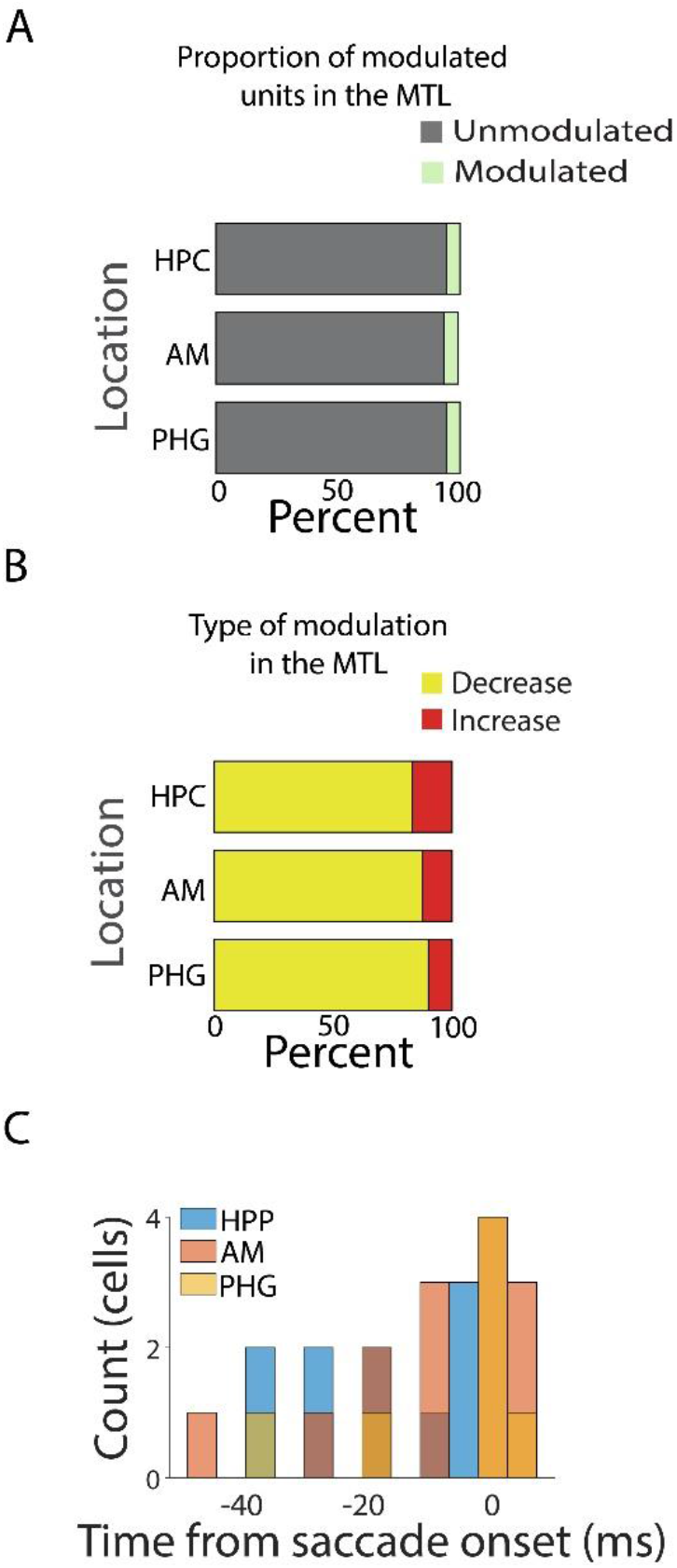
Modulatory effects are similar within different areas of the MTL. **A)** Proportion of cells modulated in the perisaccadic interval separated by the different anatomical locations; **B)** Types of modulation observed in the MTL separated by the different anatomical locations; **C)** Latencies of units modulated (decreases only) by saccade by MTL location. Note strong modulation within the saccadic onset.

**Supplementary Figure 3.**
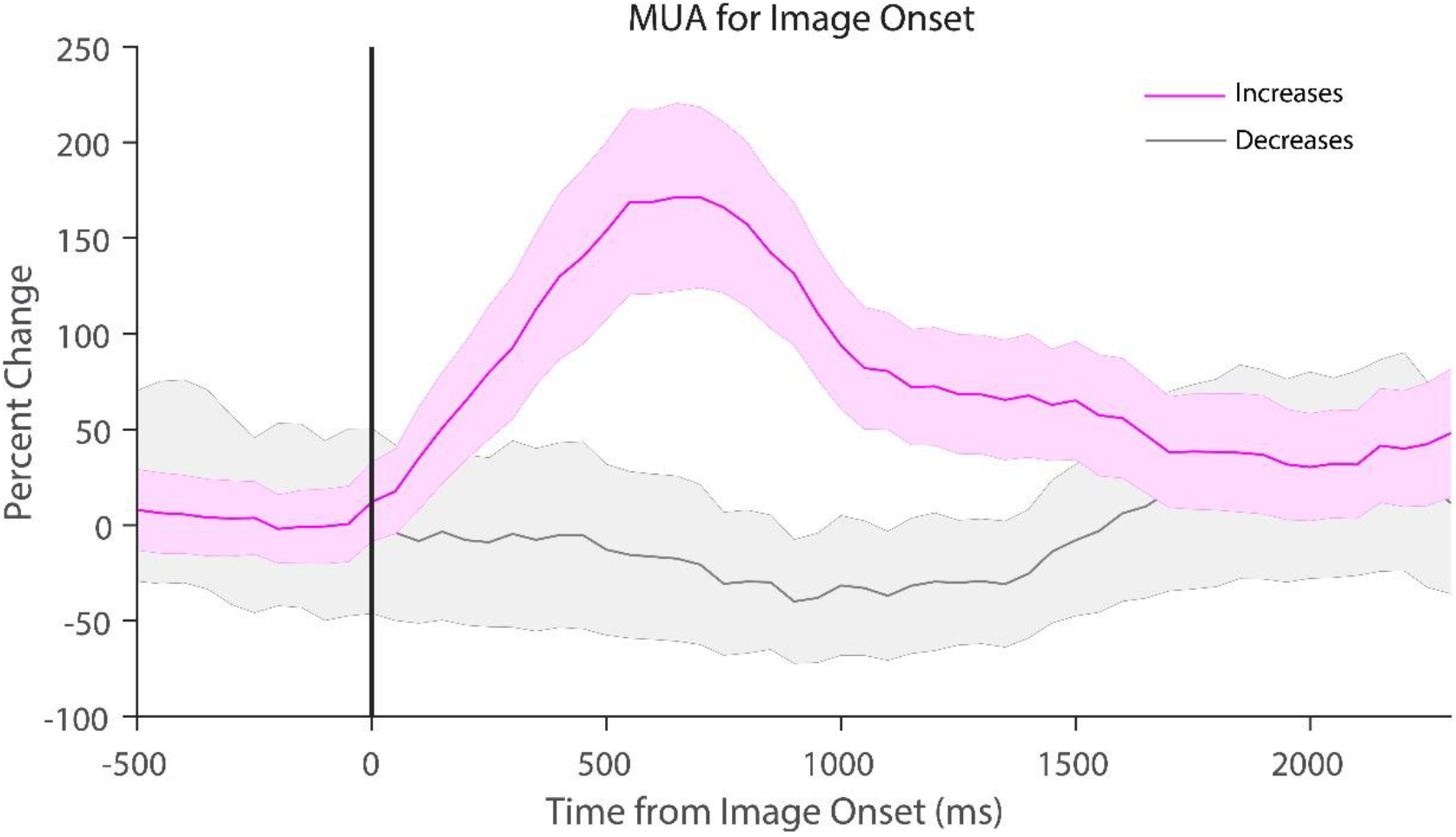
Multiple Unit Activity (MUA) of units modulated by image onset. Modulation was characterized by significant increases in firing rate assessed from 200-1700 ms. A minority of cells presented significant decreases in firing related to image onset.

**Supplementary Figure 4.**
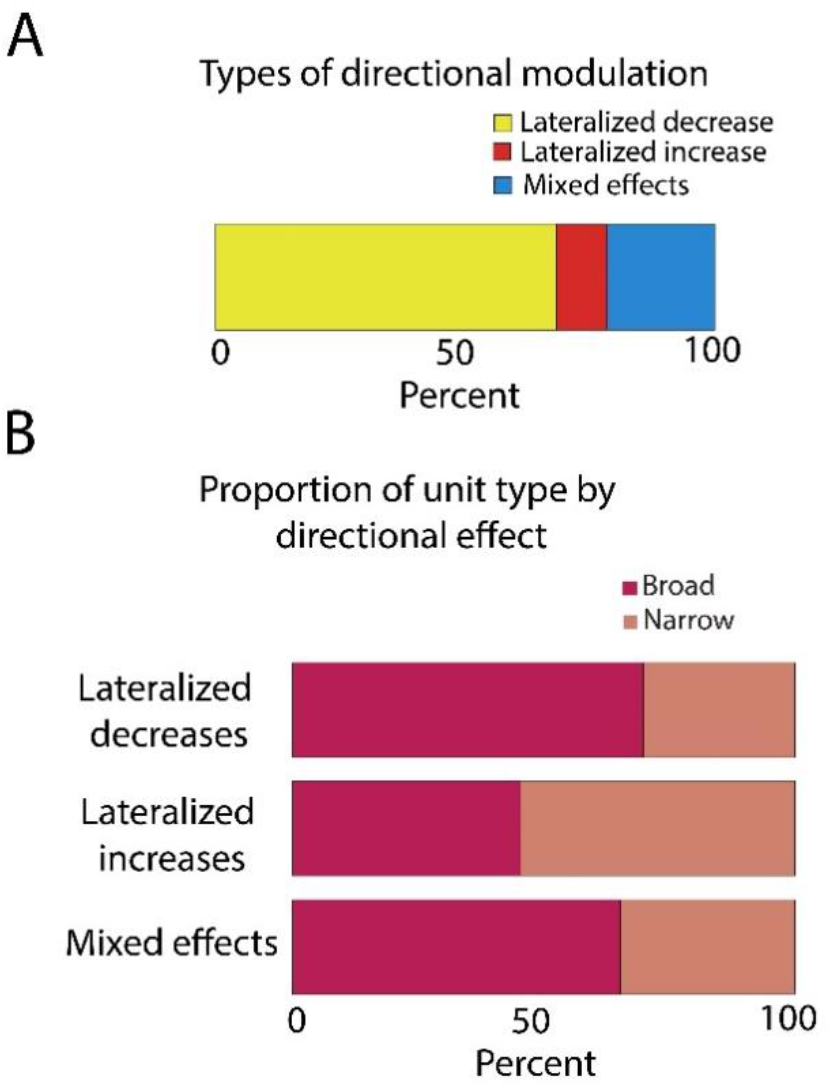
Characteristics of directional modulation in the MTL. **A)** Distribution of the different types of directional effects observed in the MTL; **B)** Proportion of ‘broad’ and ‘narrow’ spiking units separated by the directional effect observed in the MTL.

